# Arginine to glutamine mutation in olfactomedin like 3 (*OLFML3*) is a candidate for severe goniodysgenesis and glaucoma in the Border Collie dog breed

**DOI:** 10.1101/321406

**Authors:** Carys A Pugh, Lindsay L Farrell, Ailsa J Carlisle, Stephen J Bush, Violeta Trejo-Reveles, Oswald Matika, Arne de Kloet, Caitlin Walsh, Stephen C Bishop, James GD Prendergast, Jeffrey J Schoenebeck, Joe Rainger, Kim M Summers

## Abstract

Goniodysgenesis is a developmental abnormality of the anterior chamber of the eye. It is generally considered to be congenital in dogs (*Canis lupus familiaris*), and has been associated with glaucoma and blindness. Goniodysgenesis and early-onset glaucoma initially emerged in Border Collies in Australia in the late 1990s and has subsequently been found in Europe and the USA. The objective of the present study was to determine the genetic basis of goniodysgenesis in Border Collies. Clinical diagnosis was based on results of examinations by veterinary ophthalmologists of affected and unaffected dogs from eleven different countries. Genotyping using the Illumina high density canine SNP chip and whole genome sequencing were used to identify candidate genetic regions. Expression profiles and evolutionary conservation of candidate genes were assessed using public databases. Analysis of pedigree information was consistent with an autosomal recessive mode of inheritance for severe goniodysgenesis (potentially leading to glaucoma) in this breed. There was a highly significant peak of association over chromosome 17, with a *p*-value of 2 × 10^-13^. Whole genome sequences of three dogs with glaucoma, three severely affected and three unaffected dogs identified a missense variant in the olfactomedin like 3 (*OLFML3*) gene in all six affected animals. This was homozygous in all nine cases with glaucoma and 12 of 14 other severely affected animals. Of 67 reportedly unaffected animals, only one (offspring of two homozygous affected parents) was homozygous for this variant. The identification of a candidate genetic region and putative causative mutation will aid breeders to reduce the frequency of goniodysgenesis and the risk of glaucoma in the Border Collie population.

## Introduction

Companion animals including cats, dogs and horses, suffer from a range of diseases, with both genetic and environmental aetiologies. The breed barrier created by registration requirements and breeding practices seeking to maximise compliance of animals with breed specifications, results in increased levels of matings between relatives [1] and means that purebred companion animals are particularly likely to suffer from recessive genetic conditions. Understanding the genetic factors underlying these diseases is important to improve welfare within breeds and the strong linkage disequilibrium in such inbred populations means that small numbers of pedigree animals can be used to identify genetic regions of interest, making them excellent genetic models if the same disease is present in humans.

In response to the identification of causal variants, breeding strategies can be put in place to reduce the prevalence of conditions that impact on animal welfare, especially when a genetic test can be developed. However, the success of these approaches depends on understanding the mode of inheritance and level of genetic contribution to the disease, and ideally on identifying a causative mutation that can be used to develop a test for genetic status.

Primary glaucoma is a condition in which increased ocular pressure damages the retinal ganglion, leading to blindness (reviewed in [2]). In dogs (*Canis lupus familiaris*) it can be preceded by goniodysgenesis (also known as mesodermal dysgenesis), a developmental abnormality of the eye characterised by narrowing or closure of the iridocorneal angle through which the aqueous humour drains. This is associated with alterations in the structure of the pectinate ligament which traverses the drainage angle, called pectinate ligament dysplasia (PLD) [3] (more properly pectinate ligament abnormality [4]). In the Leonberger dog breed, goniodysgenesis has been associated with an increased risk of developing primary closed angle glaucoma (PCAG) [2] and in Flat-Coated Retrievers, development of glaucoma is positively and significantly related to the severity of goniodysgenesis [5]. Goniodysgenesis is generally considered to be congenital in dogs, although its subsequent progression is different among breeds. The severity is thought to increase with age in Leonbergers [2] and Flat-Coated Retrievers [6] and was shown to be strongly associated with age in Dandie Dinmont Terriers, Basset Hounds, Flat-Coated Retrievers, Hungarian Vizslas and Golden Retrievers [7, 8]. However, goniodysgenesis remained stable in Samoyeds [9] and was not associated with age in a small study of Border Collies [8].

Sudden onset glaucoma leading to blindness with bilateral loss of eyes began to be seen in young Border Collies in Australia about 15 years ago. A similar, sudden onset glaucoma in young animals subsequently emerged in the UK Border Collie population, often in dogs that were related to the affected Australian dogs. A number of dogs in the USA and Europe have now also been diagnosed with severe goniodysgenesis and glaucoma. Goniodysgenesis in Border Collies has been added to the Schedule B list of “Conditions Under Investigation” in the British Veterinary Association (BVA) Eye Scheme (http://www.bva.co.uk/Canine-Health-Schemes/Eye-scheme/) and it is highly prevalent in some Border collie lineages (see Anadune Border collie database: http://www.anadune.com/; free access but registration required). In a recent study [8], 11 of 102 Border Collies (10.8%) were reported to have moderate or severe PLD (associated with goniodysgenesis). Seven (6.9%) were mildly affected. Because of the association with a small number of popular sires and the high prevalence in some lineages, a genetic aetiology is strongly suspected in Border Collies, but the heritability and mode of inheritance of the condition is currently unknown. It is also not known why a proportion of dogs diagnosed with severe goniodysgenesis go on to develop glaucoma, but some do not, and nor do those with mild goniodysgenesis, over a life span of 15 or more years.

The aim of the current study was to perform a genome-wide analysis to find genetic regions that are associated with severe goniodysgenesis and glaucoma in the Border Collie and identify candidate mutations that might be responsible for this condition.

## Materials and Methods

Full details of the Materials and Methods, including details of publicly available data, are given in **Supplemental File S1**.

### Ethical approval

All studies were approved by the Veterinary Ethical Review Committee of the University of Edinburgh (approval number VERC 2012-8) and the Animal Ethics Committee of the University of Queensland (approval number ANFRA/MRI-UQ/565/17).

### Ascertainment of clinical status and pedigree

Goniodysgenesis was diagnosed by examination of the eyes with gonioscopy to assess the status of the drainage angle and the pectinate ligament fibres that span it. Since we had results in multiple formats from many veterinary ophthalmologists, each report was manually assessed to assign clinical status to each subject. Only individuals who had had gonioscopy analysis were included. Breeding records and pedigree data were obtained from public databases and from the breeders’ websites. Websites were accessed at various times up to February 2018. Full details including exclusion criteria are given in **Supplemental File S1**.

### Genetic analysis

DNA samples were from buccal cells collected by owners and occasionally from blood. Full details of collection and extraction procedures are given in **Supplemental File S1**. Genotyping used the Illumina 173k CanineHD Whole-Genome Genotyping Bead Chip (Illumina, San Diego, CA, USA). Results were filtered in PLINK v1.07 [10] to remove individuals that had more than 10% of missing genotypes and SNPs that had rates of genotyping < 0.95, had minor allele frequency < 0.05 in this population or deviated from the Hardy-Weinberg equilibrium in the controls with P value of less than 0.0001. Only the autosomes were considered.

Cases for the analysis were defined as severely affected and consisted of dogs that had goniodysgenesis of grades four or five under an early grading scheme, dogs whose goniodysgenesis was described as “severe”, those where the veterinary ophthalmologist report indicated PLD over more than 70% of the iridocorneal angle and those that developed glaucoma following a diagnosis of goniodysgenesis. Controls were dogs that were assessed as clear or clinically unaffected based on at least one gonioscopy examination. Previous work indicated that goniodysgenesis was not progressive in Border Collies [8] so the controls were categorised on the basis of a clear gonioscopy, regardless of age. The SNP analysis was performed in GEMMA v0.94.1 [11] using a linear mixed model approach, controlling for relatedness using a SNP-based relatedness matrix as a random effect. The significance threshold was based on a Bonferroni correction of 0.05. Additionally, a more conservative, regional approach was undertaken using sliding windows of 40 adjacent SNPs, overlapping by 20 SNPs in REACTA v0.9.7 [12] with the first two genomic principal components controlling for relatedness. Whole genome sequencing (30X coverage) of DNA from three dogs with glaucoma, three severely affected and three unaffected animals was performed using the Illumina HiSeq X platform. Library insert sizes were 450 bp and paired reads of 150 bp were sequenced. Reads were processed by standard procedures, detailed in **Supplemental File S1** and mapped to the CanFam3.1 reference genome. The significance of variants was assessed using the Ensembl Variant Effect Predictor [13].

### Transcriptomic analysis of candidate region genes

A time course of mouse eye development from embryonic day 12 to adult was examined using the FANTOM5 database [14–16] (http://fantom.gsc.riken.jp/zenbu). FANTOM5 data were also used to obtain expression levels for human eye-related samples (**Supplemental Table S1**). Transcriptomic data for the developing chicken eye were generated for another project by RNA sequencing methodology (Rainger et al, in preparation). Details are given in **Supplemental File S1**. Microarray-based expression data for a range of tissues in human and mouse were from BioGPS (http://biogps.org) as were RNA sequencing data for chicken tissues. Microarray expression data for immune cell types was derived from the Immunological Genome Project (https://www.immgen.org/).

### Analysis of canine OLFML3 gene

The region of the canine *OLFML3* gene containing the candidate causative mutation was amplified by the polymerase chain reaction and the amplicon sequenced by chain termination sequencing. Full details of primers and protocols are given in **Supplementary File S1**. Sequences for analysis of nucleotide and amino acid conservation across species were downloaded from Ensembl (http://ensembl.org) and alignment was performed using Clustal Omega (http://www.ebi.ac.uk/Tools/msa/clustalo/).

## Results

### Familial clustering of goniodysgenesis and glaucoma

Inspection of pedigrees revealed that there were three common ancestors in both the sire and dam lineages for all cases of glaucoma on the goniodysgenesis database (https://bcglaucomadatabase.synthasite.com/) (N=12) including two pairs of full siblings both affected with glaucoma, as well as additional cases not on the database (N=4). These common ancestors dated back to the late 1970s and early 1980s and were found in multiple ancestral lineages for the glaucoma cases. All the glaucoma cases had been diagnosed with goniodysgenesis and subsequently had one or both eyes removed. However, several individuals diagnosed with severe goniodysgenesis had not gone on to develop glaucoma. This included two popular sires who reached the age of 15 years without a diagnosis of glaucoma. It also included a full sibling of two dogs with glaucoma. Three dogs with severe goniodysgenesis had produced a total of six offspring with glaucoma.

### Genome wide association analysis

To identify genetic regions associated with goniodysgenesis in this breed, we performed SNP analysis on 17 severely affected cases (8 female, 9 male) and 42 unaffected dogs (26 female, 16 male) who had been passed as “clear” at veterinary ophthalmology testing. The cases all had goniodysgenesis described as “severe” (N=4), Grade 4 or 5 by the earlier grading scheme (N=1 and 5 respectively), or goniodysgenesis followed by glaucoma (N=7). A strong association of goniodysgenesis with a region of chromosome 17 was found (**Figure 1A**). The lowest *p*-value was for SNP rs22561716 at 17:51,919,221 (P=2 × 10^13^) included in a window of 40 SNPs that had a *p*-value of 3 × 10^-07^according to REACTA. SNPs with the lowest *p*-values were in a region of about 1 × 10^6^ base pairs (**Figure 1B**). Sixteen of the severely affected and glaucoma cases were homozygous for this haplotype and one case was heterozygous in this region. Within the genomic region identified by the association analysis there were a number of coding genes (**Figure 1A**), although none was known to be involved in eye development or glaucoma in animals or humans. Several had strong expression in the developing mouse eyeball (**Supplemental Figure F1**).

**Figure 1.**
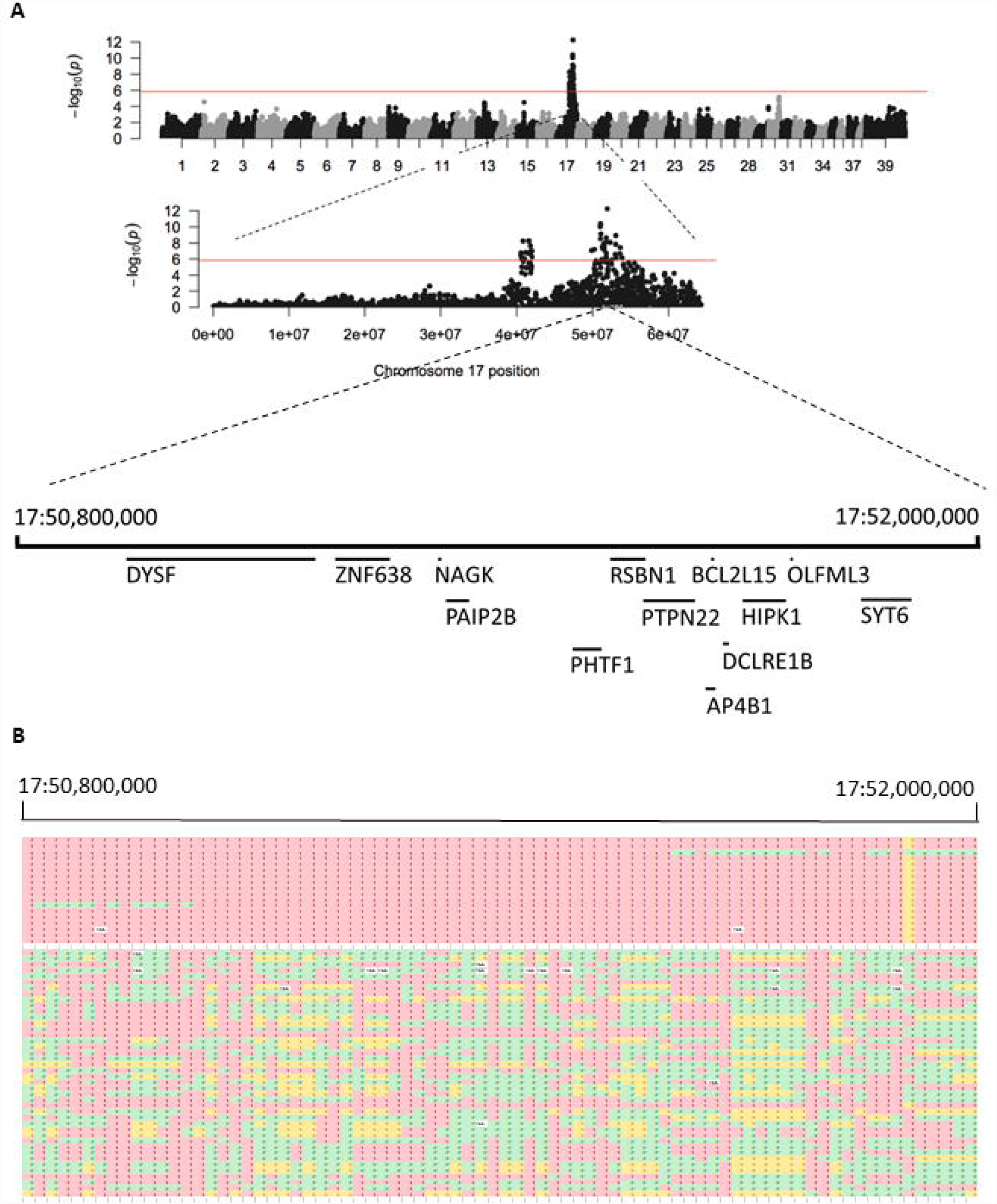
Genome wide association with severe goniodysgenesis and glaucoma **A**. Manhattan plot for the canine genome; chromosome 17 is shown in more detail and genes in the region are shown below. **B**. Homozygosity within the region of interest. Each row is a DNA sample and each column shows a SNP. Red and yellow indicate homozygosity for the major and minor allele respectively and green shows heterozygosity. The upper block shows severely affected and glaucoma animals; the lower block shows those passed as clinically unaffected at gonioscopy testing. The most significant SNP is shown as yellow in the cases (homozygous for the minor allele); one clinically unaffected animal was homozygous for this allele and one severely affected animal was heterozygous.

### Whole genome sequencing

We then obtained the whole genome sequence of this region in three glaucoma, three severely affected and three unaffected individuals, who were members of four nuclear families (**Supplemental Figure F2**). All were distantly related through multiple lineages in the extended pedigree. Relatives were chosen to highlight variants that were different between affected and unaffected family members [17]. Within the region of the lowest p-value SNP, a missense variant was found in the olfactomedin like 3 (*OLFML3*) gene, at position 17:51,786,924. The mutation was c.590G>A with the predicted change of arginine at position 197 to glutamine (p.R197Q). This arginine is in the olfactomedin (OLF) domain [18, 19] (**Figure 2A**) and was found to be highly conserved throughout mammals, birds and reptiles (**Figure 2B**). The impact predicted by the Ensembl Variant Effect Predictor [13] was given as “moderate” indicating “a non-disruptive variant that might change protein effectiveness”, consistent with the non-lethal phenotype of homozygous animals. The SIFT score was 0.55, indicating a mild effect. In humans, the mutation (c.587G>A) corresponds with rs377336789 (https://www.ncbi.nlm.nih.gov/snp/) and was seen in four out of 121,412 chromosomes with a minor allele frequency of 3.3 × 10^-5^. Genotypes are not provided for the human data, but these were presumably all heterozygous, given the minor allele frequency. SIFT score was 0.54 and PolyPhen score was 0.392, also consistent with a mild effect. The mutation was heterozygous in four of 35 Border Collies from the Dog Biomedical Variant Database Consortium (giving an allele frequency of 0.06 in this breed) but absent in 504 genotypes of other breeds. Notably, it was not present in 51 Leonbergers, three Basset Hounds, eight Golden Retrievers or one Dandie Dinmont Terrier, all breeds reported to be associated with goniodysgenesis. In addition, 350 dogs recorded as Border Collies but of unknown clinical status were genotyped by Animal Genetics (Talahassie, Florida, USA) via allele-specific PCR run on an ABI 3730 machine. There were 2 homozygotes and 29 heterozygotes among these 350 dogs, giving an allele frequency of 0.05, consistent with the smaller sample from the Consortium.

**Figure 2.**
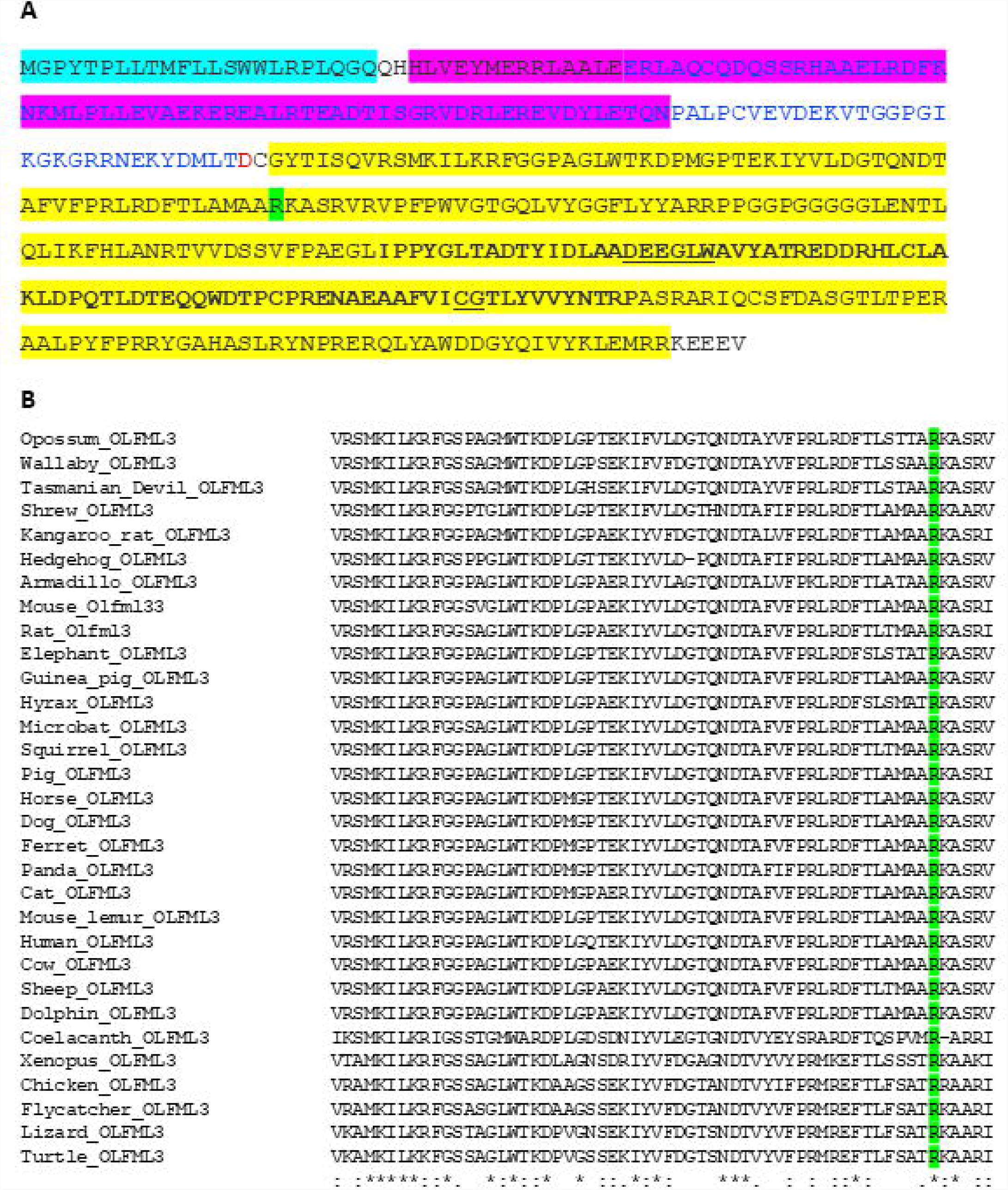
Conservation of *OLFML3* predicted amino acid sequence **A**. Amino acid sequence of the canine *OLFML3* protein. Functional domains are highlighted: blue – signal peptide; pink – coiled coil domain; yellow – olfactomedin like domain [18, 19]. Within the yellow region, bold lettering shows the core OLF region. Green shows the arginine at position 197. **B**. Conservation of arginine 197 (green) in OLFML3 of vertebrates. Arginine is found at the equivalent position in all mammal, bird and reptile species for which there is an annotated *OLFML3* gene, and in coelacanth but not in other fish. The region around this arginine is also highly conserved.

### OLFML3 genotypes of severely affected and unaffected Border Collies

The results for the mutation c.590G7#x003E;A in glaucoma, severe and unaffected dogs are shown in Table 1. The individual who was heterozygous for the region based on SNP chip was also AG at *OLFML3*. This animal had been diagnosed with severe goniodysgenesis but did not develop glaucoma over a 15-year lifespan. In addition, one unaffected dog was AA genotype. This animal was the offspring of a severely affected (grade 5) father and a mother diagnosed as affected (with no grade). Both parents were also AA. Under some testing schemes animals could pass with some degree of goniodysgenesis [2, 4] which may be the case with this dog. In general, the genotype for *OLFML3* segregated with that for rs22561716, indicating strong linkage disequilibrium between them. However, an unaffected mother and son were homozygous for G at *OLFML3* but heterozygous at rs22561716 suggesting that the ancestral haplotype was G at *OLFML3* with T at rs22561716. In addition, two unaffected individuals were homozygous for several SNPs either side of *OLFML3* but heterozygous for the *OLFML3* mutation and rs22561716 and appeared to carry a different haplotype.

**Table 1.** Genotype frequencies for *OLFML3* mutation c.590G>A in severely affected and unaffected dogs. Dogs that were included in the genome wide analysis (SNP-chip dogs) are shown separately from dogs that were only tested for *OLFML3*.

| Genotype | AA | AG | GG | TOTAL |
| --- | --- | --- | --- | --- |
| Clinical status |  |  |  |  |
| *SNP-chip dogs<sup>1</sup>* |  |  |  |  |
| **Glaucoma** | 7 | 0 | 0 | 7 |
| **Severe goniodysgenesis** | 9 | 1 | 0 | 10 |
| **Unaffected** | 1 | 19 | 22 | 42 |
| *Other dogs* |  |  |  |  |
| **Glaucoma** | 2 | 0 | 0 | 2 |
| **Severe goniodysgenesis** | 3 | 1 | 0 | 4 |
| **Unaffected** | 0 | 17 | 8 | 25 |
| **TOTALS** |  |  |  |  |
| **Glaucoma** | 9 | 0 | 0 | 9 |
| **Severe goniodysgenesis** | 12 | 2 | 0 | 14 |
| **Unaffected** | 1 | 36 | 30 | 67 |

To validate these results, we genotyped 31 additional Border Collies that had had gonioscopy investigations: two with glaucoma (both female), four severely affected (all female) and 25 unaffected (10 male, 15 female). The genotypes of these dogs are shown in Table 1 and are consistent with the results suggesting that severe goniodysgenesis/glaucoma was associated with homozygosity for the A allele of *OLFML3*.

### OLFML3 genotypes in mildly affected and uncategorised affected Border Collies

Our initial analysis compared severely or glaucoma affected dogs with dogs passed as unaffected at gonioscopy examination. Because of changes in the reporting scheme in the UK and differences among countries, there were a number of equivocal diagnoses. These were dogs where either the level of goniodysgenesis was given as mild (Grade 1 under the former scheme or stated as mild or marginal on the report; N = 9; five female, four male), moderate (Grade 2 – 3 under the old scheme, or stated as moderate or between 25 and 70% blocked on the report; N = 9; eight female, one male), or the dogs were diagnosed as affected with no description given (N = 15; nine female, six male). To examine whether *OLFML3* was involved in these cases we genotyped these 33 dogs. The results are shown in Table 2. All three genotypes were found among these cases.

**Table 2.** Genotype frequencies for *OLFML3* mutation c.590G>A in mild, moderate and uncharacterised affected dogs.

| Genotype | AA | AG | GG | TOTAL |
| --- | --- | --- | --- | --- |
| Clinical status |  |  |  |  |
| **Mild** | 3 | 3 | 3 | 9 |
| **Moderate** | 4 | 2 | 3 | 9 |
| **Affected, no grade** | 7 | 3 | 5 | 15 |
| **TOTALS** | 14 | 8 | 11 | 33 |

### Expression of OLFML3 in the eye

In expression data generated by the FANTOM5 project, the *Olfml3* gene was strongly expressed in mouse eyeball at embryonic and neonatal stages and declined in the adult sample (**Figure 3A**). FANTOM5 data showed that the gene was expressed in human lens, corneal epithelium and retina (**Figure 3B**), although the highest expression was in amniotic and placental epithelial cells and in mesenchymal stem cells (not shown). *OLFML3* was also expressed in the developing chick eye, with highest expression at embryonic day 6 (**Figure 3C**). In mouse microarray data (mouse probeset 1448475_at; http://biogps.org) expression of *OLFML3* was highest in osteoblasts undergoing calcification, but it was also found in eyecup, iris, ciliary bodies and lens (**Supplemental Figure S3A**). There was negligible expression in immune system cells, except for macrophages, osteoclasts and particularly microglia, which had expression comparable to lens and eyecup. Strong expression in microglia was also seen in the Immunological Genome data set (**Supplemental Figure S3B**). Microarray data also available through BioGPS showed strongest expression of *OLFML3* in human adipocytes, uterus and retina, the only eye tissue represented (human probeset 218162_at; http://biogps.org) (**Supplemental Figure S3C**). RNA sequencing data compiled on BioGPS revealed expression in many samples from the chicken, notably the whole embryo and connective tissues and the cornea of the adult (**Supplemental Figure 3D**). Other OLF family members were also expressed in the developing mouse eyeball with a similar pattern to *Olfml3*, peaking in the early neonatal period and declining in the adult eye (**Supplemental Figure S4**). In contrast, the Myocilin gene (encoding a related protein which is mutated in human open angle glaucoma) had negligible expression in embryo and early neonate but increased expression at neonatal day 16 and in adult eyeball (**Supplemental Figure S4**).

**Figure 3.**
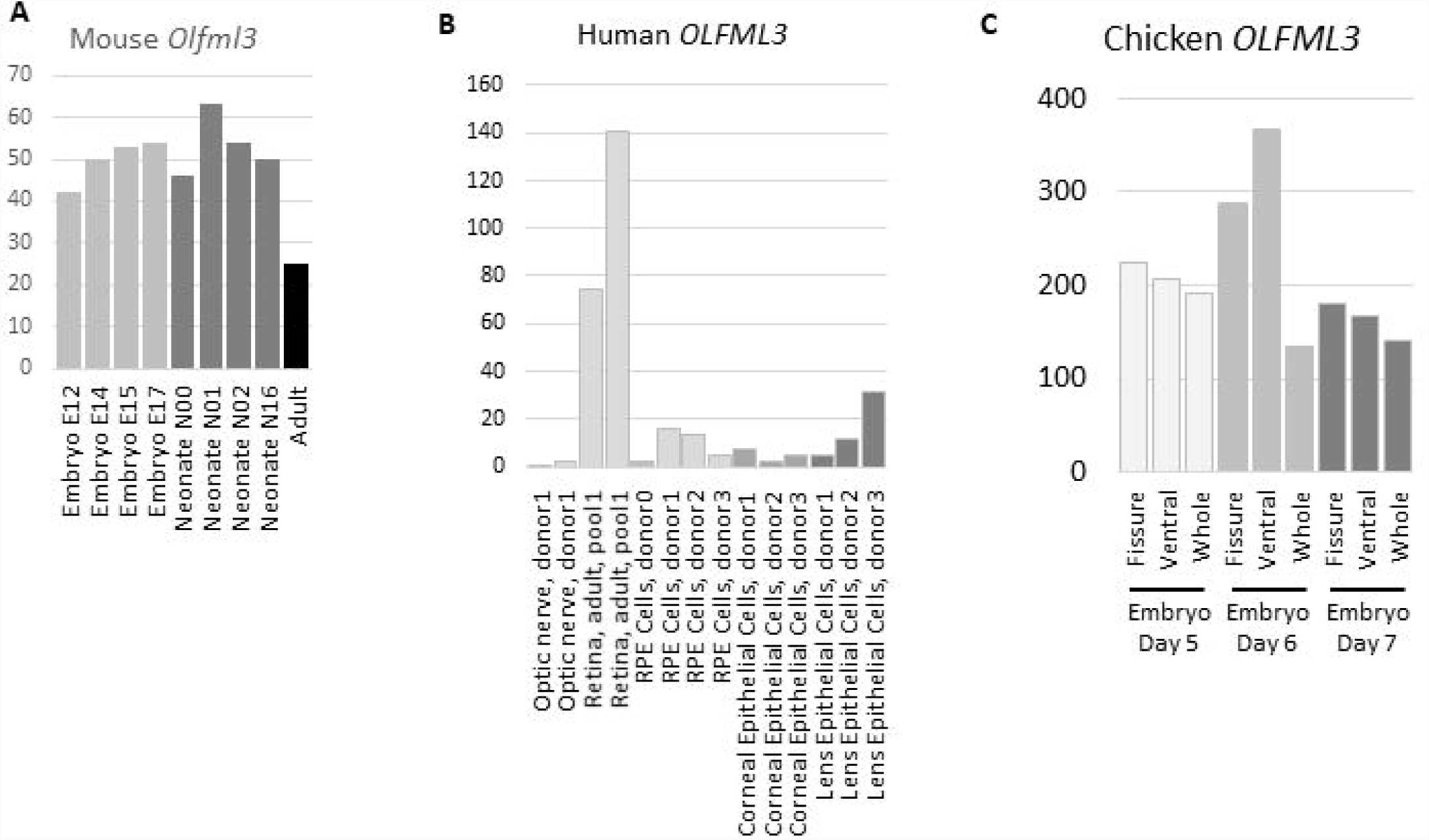
Expression of *OLFML3* genes in eye regions **A**. Expression of *Olfml3* in developing mouse eyeball. Results are based on CAGE and were downloaded from the FANTOM5 database (http://fantom.gsc.riken.jp/5/tet). Y axix shows RLE normalised values. Light grey – embryonic samples; dark grey – neonatal samples; black adult sample. **B**. Expression of *OLFML3* in human eye tissues and cells. Results are based on CAGE and were downloaded from the FANTOM5 database. Y axis shows RLE normalised values. Light grey – optic nerve and retina; mid grey – cornea; dark grey – lens. **C.** Expression of *OLFML3* in the developing chick eye. Results are based on RNA sequencing. Y axis shows transcripts per million (TPM) values. Light grey – embryonic day 5; mid grey – embryonic day 6; dark grey – embryonic day 7.

## Discussion

In a number of dog breeds development of glaucoma has been associated with the presence of goniodysgenesis. In the Flat-Coated Retriever, the likelihood of glaucoma was positively and significantly related to the severity of goniodysgenesis [5] and in the Leonberger five glaucoma-affected individuals all had a high grade of goniodysgenesis in the contralateral eye [2]. An abnormal drainage angle is considered typical of some breeds, with reports of up to 81% of individuals showing evidence of goniodysgenesis [9, 20]. However, the highest prevalence of glaucoma reported in any single breed is only 5.52% [21], indicating that severe goniodysgenesis does not necessarily proceed to glaucoma.

An increasing incidence of glaucoma in young animals has been reported in the Border Collie breed over the last 15 years. Consistent with results in other breeds, our survey of goniodysgenesis in the Border Collie found some individuals who lived entire lifetimes with apparently severe goniodysgenesis without developing glaucoma. This would suggest that although severe goniodysgenesis is important in the progression to glaucoma, it is not the sole cause. The aetiology may involve a genetic predisposition to goniodysgenesis combined with genetic, environmental or random factors influencing development of glaucoma.

The diagnosis of goniodysgenesis was made by veterinary ophthalmologists (such as members of the BVA Eye Panel in the UK or the equivalent certifying body in other countries) who performed gonioscopies to assess the status of the drainage angle and the pectinate ligament fibres that cross it. Since the diagnoses were made by a number of different clinicians over a period of years, there may be considerable noise in the phenotype. An initial eye scheme developed by the BVA in association with the Kennel Club of the United Kingdom involved assessing whether the dog had no eye abnormality (a pass), or some abnormality graded from one to five according to the extent of PLD and the width of the iridocorneal angle, in increasing levels of severity. Since there was a high degree of inter-observer variability in assigning a grade, the scheme was subsequently amended such that the dog would either pass or fail. Dogs that passed could have a drainage angle that was abnormal to some extent (“mild” goniodysgenesis; see information available at https://www.bva.co.uk/Canine-Health-Schemes/Eye-scheme/). The pass/fail scheme was expected to have a higher level of inter-observer agreement. This was confirmed in a recent study of Leonberger dogs [2] where there was a high correlation in proportion affected with goniodysgenesis between a prospective study where examinations were performed by a single specialist (2012-2014; 18% affected) and a retrospective study of BVA eye scheme certificates (2009-2014; 22% affected). In a study of PLD in Welsh Springer Spaniels, there was a good correlation between two examiners in determining unaffected eyes, although there was considerable variability between examiners in assigning the degree of PLD [4] This variability in categorising the extent of goniodysgenesis means that the classification of phenotype as “affected” or “unaffected” can result in mild PLD or angle narrowing being recorded as clinically unaffected [2, 4]. For this reason, we focussed our genome-wide analysis on severely affected dogs including several that had gone on to develop glaucoma.

Using severely affected cases and unaffected controls, we found a strong peak of association on chromosome 17, in the region of the *OLFML3* gene, where we detected a missense mutation p.R197Q. Although this gene has not been implicated in goniodysgenesis or glaucoma in humans, it is strongly expressed in tissues of the anterior segment of the human and baboon eye including lens, iris, sclera and trabecular meshwork [18, 22] and in the developing mouse eye [23]. The *Olfml3* gene was expressed throughout mouse eyeball development (**Figure 3A**) and during chick eye development (**Figure 3C**). In mouse, inactivation of the *Olfml3* gene by insertion of a *Lac* cassette resulted in viable and fertile homozygotes [23], consistent with the relatively mild selective disadvantage of dogs with goniodysgenesis. Expression has also been observed in human tissues of mesenchymal origin such as bone and adipose. Olfactomedin like 3 is an extracellular matrix protein that has been implicated in the epithelial to mesenchyme transition in cancer [24], in keeping with its mesenchymal expression and suggesting that abnormality of this protein could be associated with the formation or retention of abnormal sheets of mesenchyme in the drainage angle. Olfactomedin like 3 is also proangiogenic [25] and interacts with a member of the bone morphogenic protein (BMP) family, BMP4. Since BMPs including BMP4 are involved in eye development (see [26] for a review), this may indicate the pathway for olfactomedin like 3 action.

In addition, expression of *OLFML3* is also characteristic of microglia in humans [27, 28] and mice (http://biogps.org; http://www.immgen.org), although it is not strongly expressed by other macrophage lineages. Genetic manipulation resulting in absence of microglia is associated with loss of *Olfml3* expression in mice (R Rojo *et al*, in revision). The retina contains microglia which are important for retinal health [29, 30], and they could be one source of *OLFML3* mRNA in the canine eye. Opening of the iridocorneal angle involves considerable remodelling of the mesenchymal tissue within the angle, including thinning and extending of the pectinate ligament and opening the trabecular meshwork [31]. Microglia may contribute to this process as eye development progresses by phagocytosing apoptotic cells and debris from the remodelling. Abnormality of olfactomedin like 3 in the microglia of the eye could interfere with this function and therefore disrupt the opening of the iridocorneal angle and formation of the proper drainage channels.

Olfactomedin like 3 is a member of a large family characterised by the olfactomedin (OLF) domain [19, 32], with several members expressed in eye structures. In particular, the glucocorticoid-inducible family member *Myoc*ilin (*MYOC* gene) has been implicated in dominant open angle glaucoma in humans [33] and is strongly expressed in the trabecular meshwork of the drainage angle [34, 35]. As with *Olfml3* mutants, there is no gross phenotype in *Myoc* knockout mice [36]. Increased expression of *Myoc* mRNA resulted in a reduction in *Olfml3* mRNA [37] in a mouse model, suggesting that there is an inverse association between these two OLF proteins, consistent with the expression profiles in the developing mouse eye (**Supplemental Figure S4**) where *Olfml3* declined in the adult while *Myoc* rose. In vitro experiments indicated that *Myoc*ilin interacts with olfactomedin 3 (also known as optimedin, encoded by the OLFM3 gene) which was co-expressed with *Myocilin* in the eye [38]. Mutation in another OLF family member, *OLFM2*, has been associated with human open angle glaucoma in a small number of Japanese patients (rs779032127; p.R144Q) [39] and identified as contributing to eye development [40]. These other OLF family members may compensate for olfactomedin like 3 abnormalities, or may interact with olfactomedin like 3 in the development of the eye. Hence genetic variation in these family members may modify the impact of the *OLFML3* mutation in the Border Collies, explaining the variable phenotype of homozygotes and heterozygotes carrying the mutation.

We propose that the p.R197Q mutation in Border Collies is responsible for the presence of severe goniodysgenesis predisposing to glaucoma in homozygous animals. We have also seen some heterozygous animals with mild to relatively severe phenotypes (although none with glaucoma), but most heterozygotes have been passed as clinically unaffected at gonioscopy testing. This suggests that there may be variants at modifier loci that can compensate for the *OLFML3* mutation. In particular, the presence of many OLF proteins in the eye and the mild phenotype associated with relatively broadly expressed gene family members suggests redundancy for at least some functions. It is also possible that inadequate compensation for the putative effects of the p.R197Q mutation may be responsible for the progression from severe goniodysgenesis to glaucoma.

There were too few glaucoma cases in our study to allow identification of loci involved in progression from goniodysgenesis to glaucoma in the Border Collie and ambiguous phenotyping complicates the investigation of mild goniodysgenesis. The genetic relationship between the mild forms and the more severe forms is unknown, but the *OLFML3* variant described here does not appear to be associated with most cases of mild goniodysgenesis indicating that genetic predisposition to mild goniodysgenesis may be independent of the findings of this study. Although developing a test for the *OLFML3* mutation would allow breeders to select against homozygotes to decrease the prevalence of the risk genotype for severe goniodysgenesis, it is important not to reduce the breeding pool too much because of the risk of other recessive conditions resulting from homozygosity for alleles that are identical by descent. Therefore, we need to understand why some of our cases progressed to glaucoma at a young age and some did not before we can make recommendations to the breeders of Border Collies worldwide.

## Author contributions

KMS, CAP and LLF conceived the study. AJC, LLF and KMS curated and prepared the DNA samples. OM, SCB and LLF performed initial genetic analysis. LLF and CAP performed the SNP analysis. CAP and SJB performed the genome wide sequencing analysis. KMS and AJC designed, performed and analysed the *OLFML3* genotyping. JJS analysed the data from the Dog Genome Consortium. AdeK and CW analysed the Animal Genetics samples. KMS analysed the public gene expression data. JR and VT-R created and analysed the chicken eye data. KMS, LLF and CAP drafted the text and all authors read and edited the final manuscript.

## Acknowledgements

We are grateful to Dogs Trust (UK) for a Bateson Canine Welfare Grant for this project and to the Pastoral Breeds Health Foundation (UK) for pump priming funding. The Roslin Institute is supported by core funding from the Biotechnology and Biological Sciences Research Council (UK) [Grant numbers BB/J004235/1, BB/J004316/1, BB/P013732/1]. JR is funded by Fight for Sight (UK) [Grant number1590/1591]. VT-R was funded by a CONACYT (Mexico) international studentship. KMS is supported by the Mater Foundation (Brisbane, Australia). We would like to thank all the Border Collie owners and breeders who have contributed samples and information to this investigation and the Pastoral Breeds Health Foundation for continuing support and encouragement during the course of this project. We are grateful to Dr Alan Wilton (now deceased) and Dr Barbara Zangerl for initiating and supporting the project.

## Supplemental Material

**Supplemental Table S1**. FANTOM5 samples used in analysis of gene expression. Details can be found at fantom.gsc.riken.jp/5/sstar/.

**Supplemental Figure S1**. Expression of genes in the candidate region in the developing mouse eye. Upper panel shows the canine genes in the region. Lower panel shows expression of the genes in the developing mouse eyeball. Y axis shows RLE normalised expression levels. Data taken from FANTOM5 (see **Supplemental Table S1**). Light grey – embryonic; dark grey – neonatal; black – adult.

**Supplemental Figure S2**. Pedigrees of animals used for whole genome sequencing. Grey fill indicates severe goniodysgenesis; black fill indicates glaucoma; white fill indicates unaffected or unknown status. Double line indicates a first cousin mating; all glaucoma individuals can be traced back to the same three distant common ancestors on both sides. Animals with a BC ID were sequenced.

**Supplemental Figure S3.** Expression of olfactomedin like 3 gene in cells and tissues

A. Expression in mouse tissues and cell lines (http://biogps.org; microarray probeset 1448475_at). Y axis shows relative intensity of expression (arbitrary fluorescence units).

B. Expression in mouse myeloid cells (http://www.immgen.org, dataset “MFs, Monocytes, Neutrophils” microarray probeset 10500808). Y axis shows relative intensity of expression (arbitrary fluorescence units).

C. Expression in human tissues and cell lines (http://biogps.org; microarray probeset 218162_at). Y axis shows relative intensity of expression (arbitrary fluorescence units).

D. Expression in chicken tissues (RNA sequencing data curated from public databases and analysed using Kallisto, available at http://biogps.org/dataset/BDS_00031/chicken-atlas/; Bush *et al*, in revision). Y axis shows tags per million (TPM).

In A, C and D, blue – nervous tissues; dark green □ connective tissues; light green □ digestive tract and renal; yellow – reproductive; orange – immune; red – ocular.

**Supplemental Figure S4**. Expression of olfactomedin family genes in the developing mouse eye (data from FANTOM5; http://fantom.gsc.riken.jp/zenbu/). Y axis shows RLE normalised expression levels based on CAGE analysis. Light grey – embryonic; dark grey – neonatal; black – adult.

